# Nanoscale imaging approaches to quantifying the electrical properties of pathogenic bacteria

**DOI:** 10.1101/142315

**Authors:** Ryan Berthelot, Suresh Neethirajan

## Abstract

Biofilms are natural, resilient films formed when microorganisms adhere to a surface and form a complex three-dimensional structure that allows them to persist in a wide variety of environments. Readily forming in hospitals and on medical equipment, biofilms are frequent causes of infections and their subsequent complications. Due to the complexity of these structures, systematically studying individual bacterial cells and their interactions with their surrounding environment will provide a deeper understanding of the processes occurring within the biofilm as whole versus bulk population based methods that do not differentiate individual cells or species. Methods based on atomic force microscopy (AFM) are particularly suited to the study of individual cells, but are underutilized for the study of bacterial electrical properties. The ability of electrical currents to impair bacterial attachment is well documented, but to utilize electrical current as an effective antibacterial treatment, it is important to understand the electrical properties of bacteria. Therefore, we used AFM, Kelvin probe force microscopy, and ResiScope to measure the surface potential and conductance of *Psuedomonas aeruginosa* and methicillin resistance *Staphylococcus aureus* (MRSA) on gold and stainless steel. This is the first study to directly measure the electrical resistance of single bacterial cells using ResiScope. Our goal was to develop a framework for measuring biological molecules using conductive atomic force microscopy. We found that the average resistance for *P. aeruginosa* was 135.4±25.04 GΩ, while MRSA had an average of 173.4±16.28 GΩ. Using KPFM, the surface potential of MRSA shifted from −0.304 V to 0.153 V and from −0.280 V to 0.172 V for *P. aeruginosa* on gold versus stainless steel substrates, respectively. In an attempt to identify a potential charge carrier, peptidoglycan was also measured with the ResiScope module and shown to have a resistance of 105 GΩ.

## 1. Introduction

The World Health Organization (WHO) has deemed antimicrobial resistance a cardinal threat facing humanity [1]. A report from the Centre for Disease Control (CDC), published in 2015 underscores this threat, estimating that approximately one in every 100 individuals infected with drug resistant microbes will perish from such an infection [2]. This has caused a great deal of interest in the development of alternate preventative and treatment methods effective against resistance species of pathogenic bacteria, such as *Pseudomonas aeruginosa* and Methicillin-resistant *Staphylococcus aureus* (MRSA).

Not only do bacteria gain resistance due to the heavy and widespread use of antimicrobials in medicine, cleaning products, soaps, and sanitizers [1], but specialized communities of bacteria cells, known as biofilms, are associated with inherent antimicrobial resistance and recalcitrance to removal. Biofilms can easily proliferate within the human body, especially at wound sites where the protective epithelium is compromised. Any biofilm infection, if not cleared, can be life-threatening. Biofilms are initialized when bacteria adhere to a surface and begin to accumulate and produce an extracellular matrix, which enmeshes and protects these organisms, while increasing their adhesion strength. While antimicrobials are effective at eliminating the outer layers of bacteria near the surface of the biofilm, the bottom layer, which is tightly bound to the substrate and protected by the surrounding biomass, can persist and continue to proliferate [3,4].

MRSA and *P. aeruginosa* are opportunistic pathogens that form biofilms and are commonly associated with nosocomial infections [5]. *P. aeruginosa* is a Gram-negative, rod-shaped bacterium that is highly proficient in forming biofilms. The ability of *P. aeruginosa* to attach and form biofilms on a range of metallic and non-organics surfaces, has caused serious issues in hospitals (e.g. catheter infections in immunocompromised individuals) [6]. *P. aeruginosa* is also responsible for complications in the treatment of illnesses that affect the respiratory system, such as pneumonia or cystic fibrosis. Once inside the lungs, *P. aeruginosa* excretes a “slime” matrix composed of various biological components that can affect cilia movement and cause inflammation within the lungs, thus causing breathing complications [6]. This biofilm forming ability makes *P. aeruginosa* especially dangerous when mixed with other bacteria, as biofilms can turn a surface that is non-ideal for bacterial attachment into a surface to which other bacteria, such as MRSA, will preferentially bind [5]. A single microbe attaching to a surface, can initiate a chain reaction, creating an environment and surface more prone to infection of additional foreign, pathogenic species, such as other bacterial strains, fungi and viruses [7,8]. MRSA is Gram-positive cocci that once attached to a surface, secretes adhesins and toxins, facilitating attachment, all while damaging the surrounding human cells and tissue [9]. Therefore, these two organisms are particularly threatening to human health and require unique strategies to prevent, manage, and treat infection.

In terms of creating antibacterial treatments that target bacterial biofilms, electricity has shown great potential in disrupting bacterial adherence. The utilization of electricity as an antimicrobial therapy has been investigated not only because it can reduce bacterial attachment, but it may also accelerate the wound healing process, serving two essential functions in wound healing therapy [10–12]. Cathodic current across conductive surfaces can reduce attachment by up to 80%, potentially leading to the complete detachment of adherent bacteria [10]. Furthermore, alternating current in wound healing applications on guinea pigs, speeds healing times when directly applied to incised regions on the skin [13]. Therefore, researchers have begun creating devices that allow for the application of electricity to a wound site. For example, Park et al. created a polyester material coated with small, alternating Zn and Ag dots placed in an array across the surface. The difference in electrical potentials between the dots caused microcurrents to be generated across the wound site, affecting bacterial migration [14]. This is based on the principle that under the influence of an electrical current, the motility of bacteria, such as *P. aeruginosa* and *Escherichia coli* is impaired or guided in a specific direction via electrotaxis [15]. Although the electrical inhibition of bacterial attachment has been well documented, we do not fully understand the electrical properties of bacteria and how this influences bacterial behavior.

New state-of-the-art methods are being applied to characterize the surfaces charges of bacteria and the impact of current on bacteria. Among these, AFM is a nanoscale imaging technique that measures the topography of a surface. It has tremendous advantages over other nano-imaging techniques such as scanning electron microscopy, in that it does not require drastic changes in the environment (e.g. changes in pressure or temperature) to successfully image a sample [16]. ResiScope is a module for AFM, in the category of conductive AFM (CAFM) that simultaneously measures the current and resistance of a sample [17]. Similarly, Kelvin probe force microscopy (KPFM) is a conductive mode of AFM that simultaneously measures the contact potential difference between the tip and the sample. This value is a measure of the surface potential [18]. This method of simultaneous acquisition provides a tremendous advantage in relating the electrical properties of a sample to its physical structure. In this study, we utilized AFM, KPFM and the ResiScope to simultaneously acquire topography, surface potential, and current to measure the electrical resistance and surface potential of single MRSA and *P. aeruginosa* cells. Our goal was to develop a framework for measuring biological molecules using conductive atomic force microscopy (AFM), measures that will help us to understand how electrical current impacts pathogenic bacteria.

## 2. Methods

### 2.1 Bacterial strains and culture conditions

Strains of *Pseudomonas aeruginosa* and Methicillin-resistant *Staphylococcus aureus* (MRSA) were obtained from Dr. Scott Weese and Joyce Rousseau of the Pathobiology Department at the University of Guelph. Strains were streaked and cultured on 5% blood agar plates and grown at 37°C for 24 hours before use. Individual colonies were chosen and transferred into 6 mL of tryptic soy broth (TSB) solution to culture at 37°C in a shake incubator rotated at 200 rpm for 24 hours. One mL of the bacterial broth was then transferred to 1.5 mL microfuge containers and centrifuged at 1000 ×*g* for 5 minutes. The excess liquid was then decanted, and the remaining cells were resuspended into 1 mL of deionized water. This rinsing processes was repeated twice.

### 2.2 Sample preparation for AFM/KPFM/ResiScope imaging

Gold covered stainless steel discs, stainless steel discs, and sputter-coated gold mica substrates (~100 nm coating) were used in these experiments as conductive interfaces. Clean, gold and stainless steel discs were sonicated for 1 minute in de-ionized water to remove excess waste from the surface. Substrates were then rinsed with 1 mL of deionized water and left to dry for 4 hours. Four-hundred μL of the resuspended, rinsed bacterial solution was placed onto the stainless steel and the Au-coated mica, while 250 μL were deposited onto the gold discs (amount chosen to maximize surface coverage for each sample). Substrates were then incubated at room temperature for 20 minutes, allowing for bacterial attachment. This length of time produces highly dispersed populations for single cell imaging on the atomic force microscope. The excess solution on the substrates was then gently rinsed with 1 mL of deionized water using a micropipette. Substrates were allowed to dry overnight before imaging.

### 2.3 Preparation of Au-Coated mica

Mica sheets were cut into approximately 1.8 × 1.8 cm squares using sharp scissors. Individual sheets were cleaved to ensure a clean, flat surface with no flaking. Mica sheets were then sputter-coated under vacuum with ~100 nm of gold. This thickness allows for high current application in measurements without the degradation of the thin metal film.

### 2.4 Imaging AFM/KPFM/Resiscope

All AFM imaging was done on the Agilent 5500 series AFM. The ResiScope module from Concept Scientific Instruments was used to make current and resistance measurements.

The conversion of resistance imaging units in Volts to Ohms is given by *R* = 10^(*V*+2)^. All ResiScope images were taken with a potential of 0.7 V applied to the conductive substrate. Conductivity imaging was done using conductive PtIr-coated Si tips of 75 kHz, 2N/m and 225 nm in length. KPFM measurements were taken using Mikromasch Pt coated cantilevers of 75 kHz, 2N/m and 240 nm. Measured Cells were chosen based upon the lack of surrounding, noticeable contamination (e.g. EPS), as well as their total exposed circumference. Single cells were targeted for measurement in this experiment, leaving roughly at least half of the total cell circumference exposed.

### 2.5 Raman Spectroscopy

All Raman measurements were done on the Advantage series near-infrared Raman spectrometer by DeltaNu. A wavelength of 785 nm was used to collect spectra. Three centrifuged, rinsed quantities of MRSA and *P. aeruginosa* were combined and resuspended in deionized water to ensure a high concentration of cells. Twenty μL of rinsed bacterial solution was then deposited onto gold SERS substrates from Ocean Optics and left to dry for a period of three days. Purified peptidoglycan samples from *Staphylococcus aureus* were bought from Sigma-Aldrich. A 4 mg/mL solution of peptidoglycan was left to settle for 1 minute. Twenty-five μL of the separated and suspended peptidoglycan sample was then pipetted onto the SERS substrate and left to dry for three days.

## 3. Results

### 3.1 Bacterial resistivity and potential measurements

Gold and stainless steel substrates were chosen for their conductive properties and medical relevance. Conductive measurements were made on uncolonized bare and gold-coated stainless steel and on the same surfaces after *P. aeruginosa* and MRSA were allowed to adhere. Due to the roughness of the stainless steel, along with its increased reactivity (versus gold), we were unsuccessful in taking measurements on this surface using the ResiScope. However, we were able to acquire CAFM images of MRSA and *P. aeruginosa* on gold-coated surfaces (Figure 1). Table 1 shows the average resistance for bacterial cells adhered to the surface. The average resistance for *P. aeruginosa* was 135±25 GΩ, while MRSA had an average of 173±16 GΩ. The two values were not significantly different from one another. However, we may see differences with a larger sample size, as the average is based on resistance values from an average of four cells. These measurements are in alignment with those obtained in another study that quantified the resistance of *E. coli* cells suspended across two, conductive gold interdigitated fingers; results resistance values were ~396 GΩ [19].

**Figure 1.**
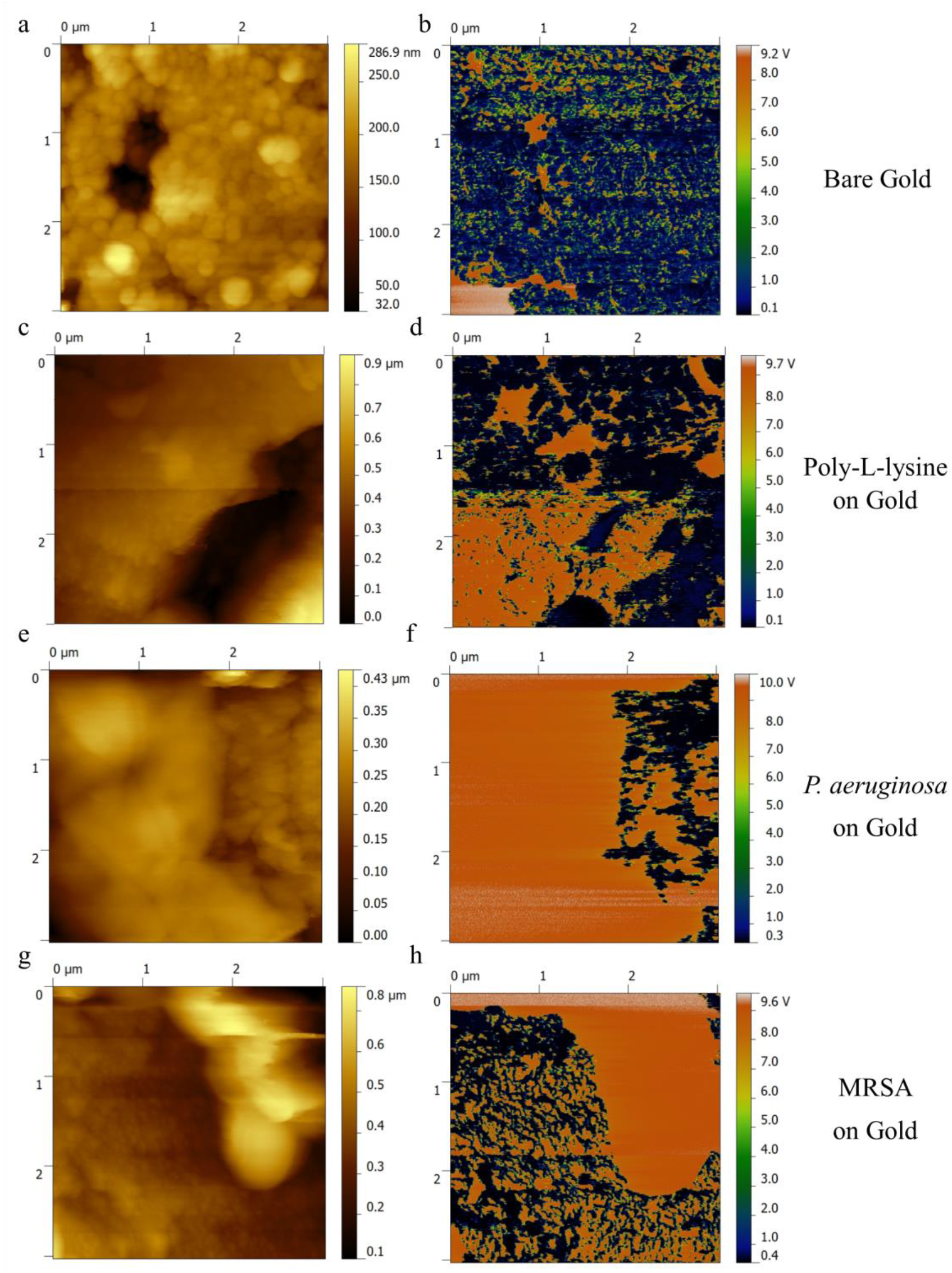
Topography and ResiScope resistance images of (a, b) bare gold, (c, d) gold-coated with poly-L-lysine, (e, f) *P. aeruginosa* and (g, h) MRSA on gold. Application of poly-L-lysine causes a slight increase in highly resistant artifacts on the surface of the gold, but the surface still remains mostly highly conductive.

**Table 1.** Average resistance and surface potential values for *P. aeruginosa* and MRSA on gold and stainless steel. Increased surface roughness and tip-sample interactions rendered ResiScope unable to retrieve viable measurements of resistance for stainless steel.

|  | Average Resistance: Au Substrate (GΩ) | Average Resistance: SS Substrate (GΩ) | Surface Potential on Gold Substrate (V) | Surface Potential on SS Substrate (V) |
| --- | --- | --- | --- | --- |
| MRSA | 173.4±16.28 | --- | -0.304±0.05 | 0.153±0.06 |
| *P. aeruginosa* | 135.4±25.04 | --- | -0.280±0.09 | 0.172±0.02 |

We also acquired KPFM measurements of MRSA and *P. aeruginosa* on stainless steel and gold-coated substrates to gain a better understanding of the relationship between surface potential and potential charge transfer (Figure 2, Table 1). Both of the bacterial cell surface potentials switched from negative to positive between the gold and stainless steel substrates. MRSA had a surface potential of −0.304 V for gold and 0.153 V for stainless steel, and *P. aeruginosa* −0.280 V for gold and 0.172 V for stainless steel. A similar effect was reported by Birkenhauer et al., where the surface potential of MRSA cells switched from positive on stainless steel to negative on gold. This suggests that surface potential varies across bacterial strains and is influenced by the surface to which the cells bind [5]. Finally, in an attempt to measure charge transfer, a 500 × 500 nm close-up resistance image of *P. aeruginosa* on gold was acquired (Figure 3). KPFM close-ups emphasize the difference in surface potential at the cell-gold interface.

**Figure 2.**
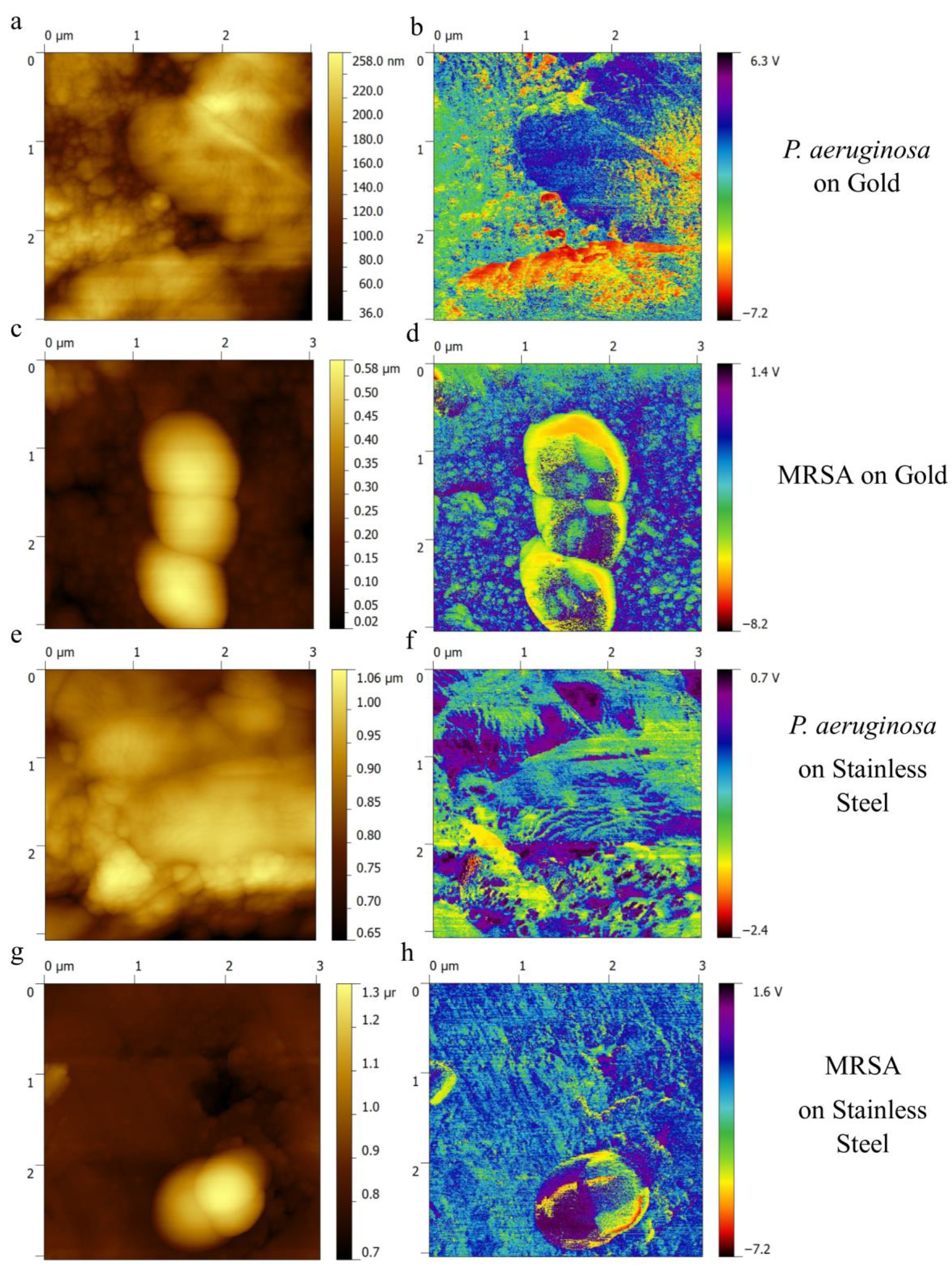
Topography and KPFM (surface potential) images of (a, b) *P. aeruginosa* on gold, (c, d) MRSA on gold, (e, f) *P. aeruginosa* on stainless steel and (g, h) MRSA on stainless steel. Surface potential changes from negative for *P. aeruginosa* and MRSA on gold to positive on stainless steel for both bacteria.

**Figure 3.**
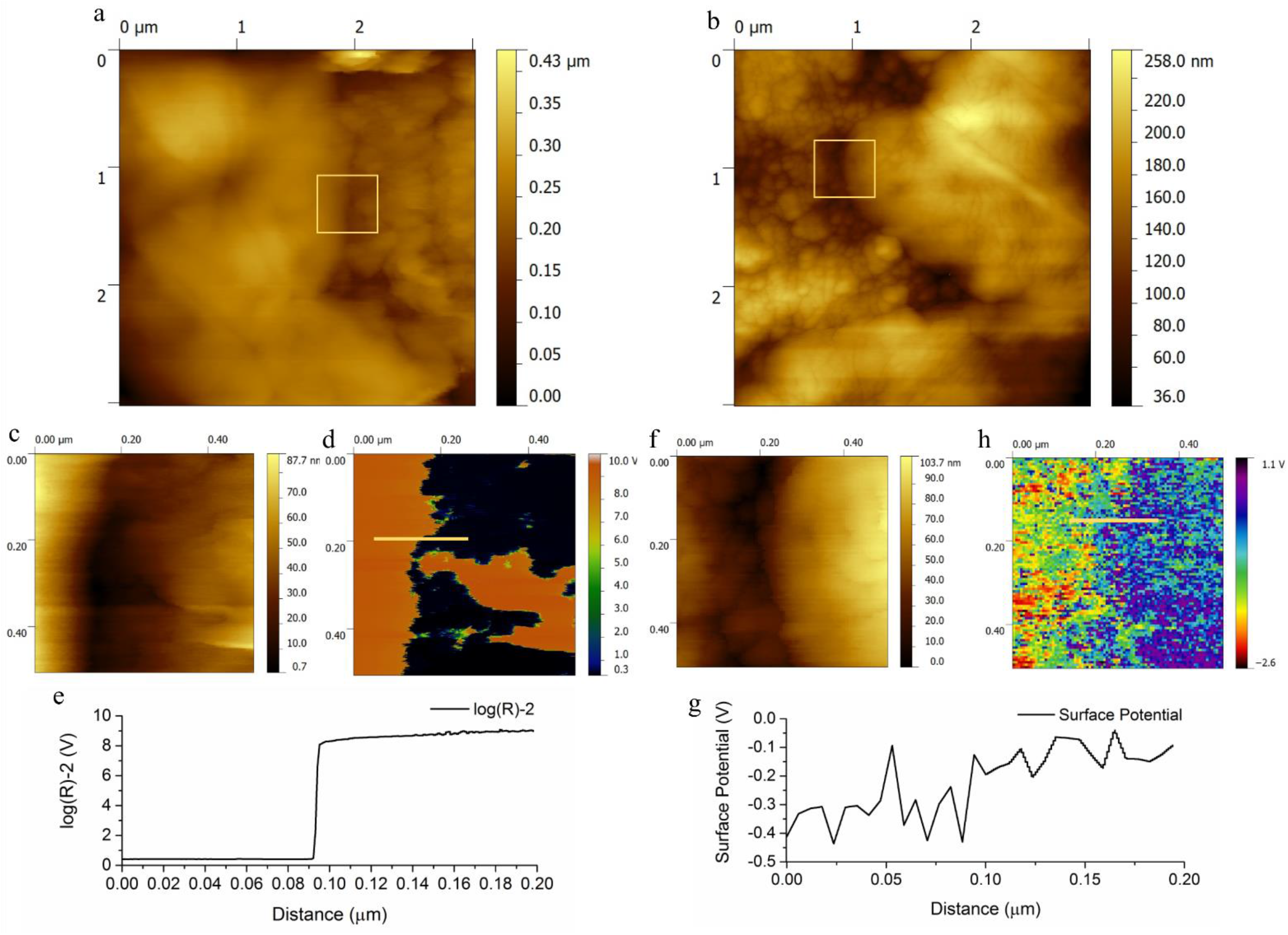
Original 3 × 3 μm zoomed out images of (a, b) *P. aeruginosa* on gold substrates. Zoomed 500 × 500 nm topographical scan (c) of *P. aeruginosa* membrane interface with gold substrate and the corresponding resistance image (d). A cross section of the resistance image (e), showing a steep change in resistance. Cropped 500 x 500 nm topographical scan (f) of *P. aeruginosa* membrane interface with gold and the corresponding surface potential image (g). Line measurement (h) of surface potential step from gold to bacterial surface.

### 3.2 The role of peptidoglycan in potential measures

We went on to investigate which bacterial substances might be involved in forming the conductive path shown in Figure 1. In solution, bacteria are conductive due to the functions of their ion channels. However, their outer membranes are largely composed of polysaccharides and lipids, which are non-conductive. Based on the structural properties of the membranes in MRSA and *P. aeruginosa*, the bulk of the electrical measurements would be made while in contact with the outer peptidoglycan layer. Therefore, we dried a 4 mg/mL of peptidoglycan on gold coated mica for 24 hours at 37°C to image the electrically conductive properties of peptidoglycan to determine if its resistive properties correlate with images of bacteria taken earlier. We found micrometer sized islands of peptidoglycan had formed, with a height of approximately 40 nm in height (Figure 4). To confirm that the purified peptidoglycan was indeed peptidoglycan, Raman spectroscopy was used, and the results were correlated with scans of *P. aeruginosa* and MRSA that show it is present in both samples (Figure 5). The peak at ~730 nm^−1^ is attributed to the stretching of the glycosidic rings in the NAG and NAM molecules present in the peptidoglycan layer surrounding the cells [19–20]. Relative intensity measures show a smaller peak for *P. aeruginosa*, most likely due to the thinner layer of peptidoglycan and its partial obstruction by lipopolysaccharides. This confirms that the molecule being measured on the surface is in fact peptidoglycan. CAFM measurements of peptidoglycan on the gold sputter coated mica show an average resistance value of 9.021 V or 105 GΩ (Figure 6). This resistance value will change depending of the thickness of the peptidoglycan sample.

**Figure 4.**
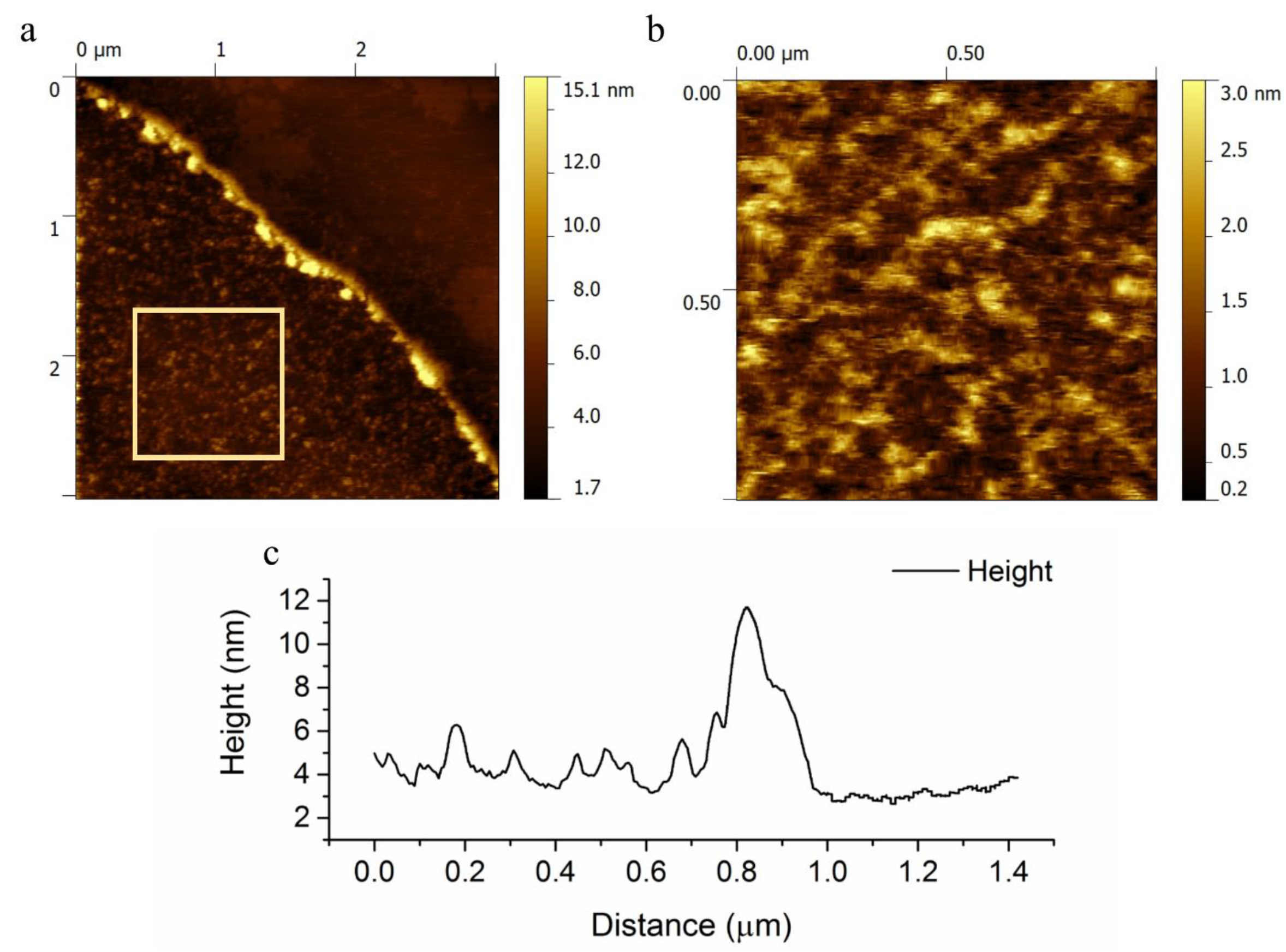
Topography images of peptidoglycan particle (a) on Au coated mica surface and (b) zoomed in 3 × 3 μm scan of particle edge. Line scan of (c) peptidoglycan particle at the Au coated mica interface, showing a height of ~40 nm.

**Figure 5.**
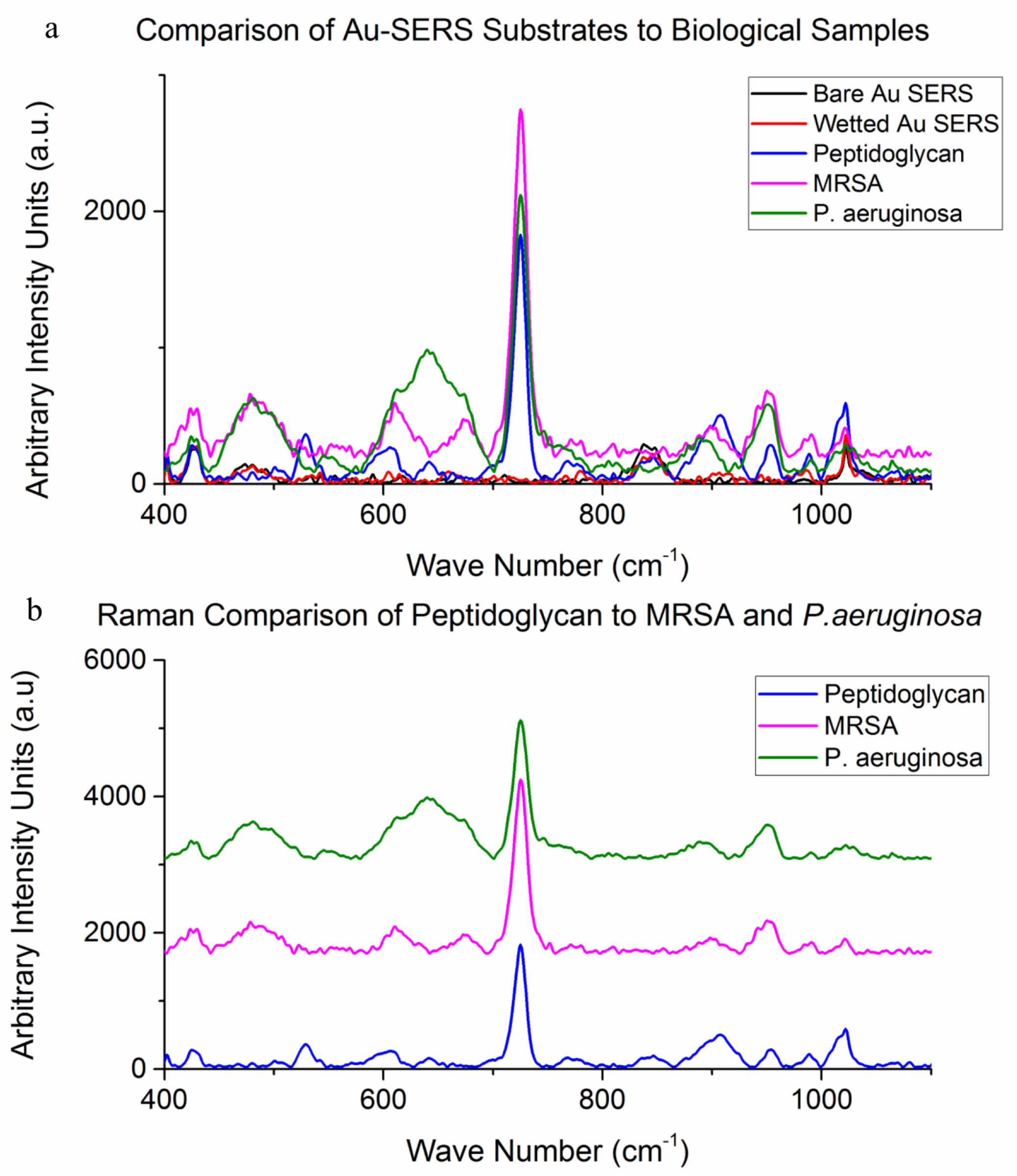
Raman spectra of (a) peptidoglycan in comparison with gold SERS substrate, *P. aeruginosa* and MRSA. Pristine and wetted-then-dried SERS substrates reference spectra show no correlation with peaks of interest. Normalized (b) spectra of peptidoglycan, *P. aeruginosa* and MRSA showing a ~730 nm^−1^ peak corresponding to glucose rings NAG and NAM molecules, which reside in the peptidoglycan.

**Figure 6.**
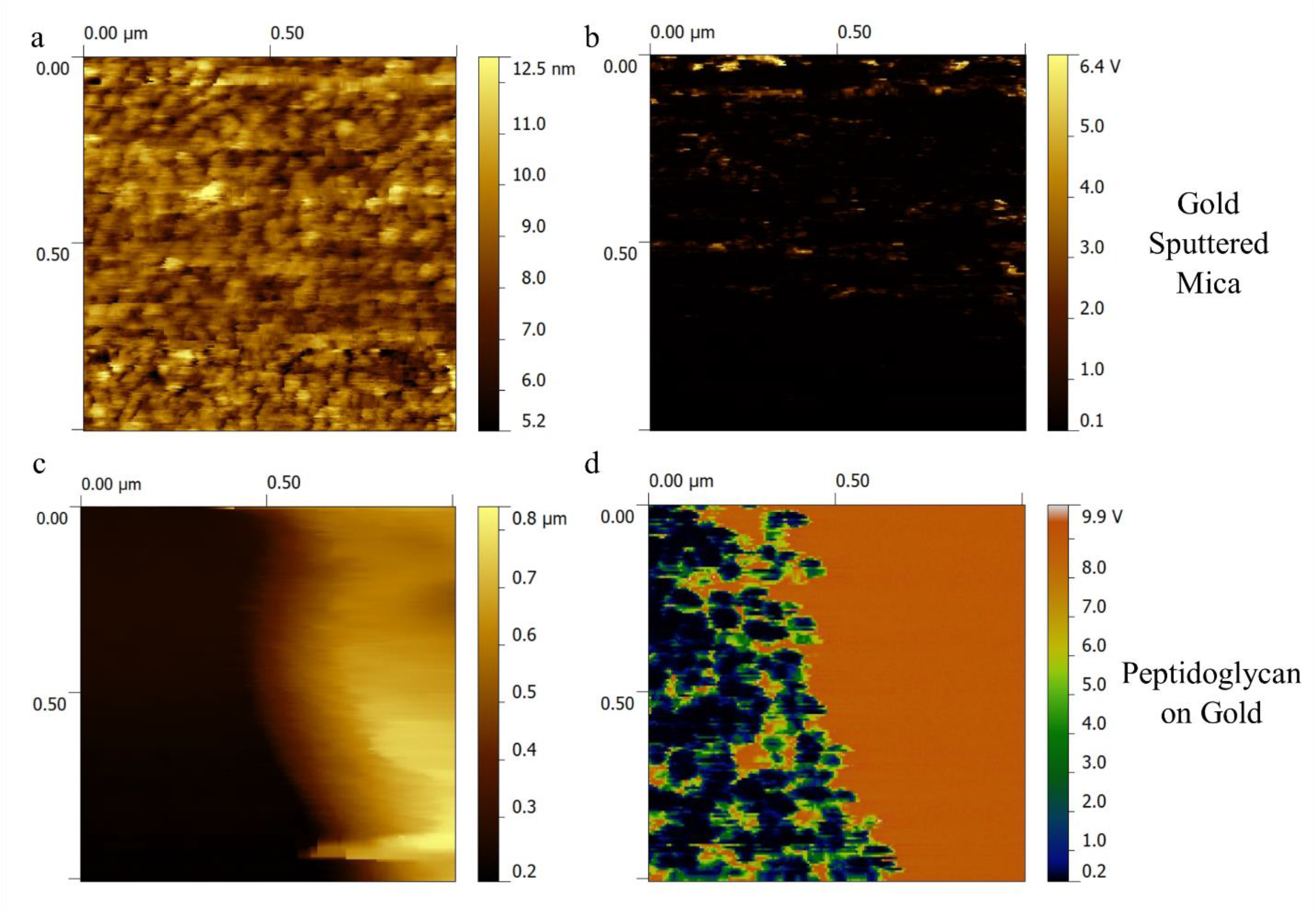
Topography and resistance 1 × 1 μm images of (a,b) bare gold and (c,d) peptidoglycan. This peptidoglycan image is from the same peptidoglycan particle as in Figure 4. The average resistance value for peptidoglycan particle on gold surface is 103 GΩ.

## 4. Discussions

AFM, KPFM, and ResiScope are excellent tools for studying the electrical potential of bacteria, as demonstrated by our results. However, the conditions must be carefully controlled to ensure ideal measurement. Controlling the quality of a ResiScope image can be accomplished by varying the force setpoint and the potential applied on the substrate. By increasing the setpoint and increasing the tip force exerted upon the sample, increased contact improves the consistency of the conductance measurements. However, if the force is raised too high, it can wear the conductive coating off the tip. Another option to produce a better electrical signal is to increase the potential applied to the substrate, thus raising the flow of current from the substrate to the tip. However, raising the voltage too high can increase the reactivity between the tip and the sample, leading to degradation of the thin metal. For these reasons, the choice of tip becomes critical [17]. Originally, we used the same platinum (Pt) tips for CAFM and KPFM. However, the Pt tips eroded rapidly, wearing away the metal coating. For these reasons, PtIr tips were used, which increased the signal response by providing a more stable flow of electrons [21]. PtIr contact mode tips with a low spring constant of 0.1 N/m are superior to Pt tips, but still generate a significant amount of noise. After testing PtIr tips with a midrange spring constant of 3 N/m and 75 kHz, we found these were ideal for our task. The higher spring constant allowed for more consistent contact between the tip and the surface. This combination gave the best and most consistent measurements for both biological and test samples. Our experiences underscore the importance of carefully selecting best settings and tips for CAFM analysis; we found that until a method is established for a particular sample type, this is likely to be a trial and error process. Our work establishes a set a parameters that can be used as a starting point for subsequent studies of a similar nature.

As for CAFM, we found that there are a variety of parameters that influence the success of ResiScope analysis. The version of ResiScope currently used by our laboratory is limited to contact mode. Contact mode, especially when used for inorganic samples, allows for better electrical contact between the tip and the substrate, which leads to more stable measurements. However, there are some shortcomings to using contact mode for these studies. First, it can easily dislodge a sample from the surface if it is weakly bound. Second, the high friction force of the cantilever pressing onto the metallic surface quickly erodes the thin film of conductive material deposited onto the Si tips [16]. There is a potential for combining KPFM and ResiScope images to study charge transfer, as a result of differences in the localized potential between the cell and the substrate. Our results give a reasonable starting point for such studies, but additional optimization is necessary to improve image quality for our application.

Our results and experiences demonstrate that there are a variety of options for enhancing and optimizing the measurement of bacterial electrical properties using CAFM. The experiments in this study provide a framework for future analysis and highlight several pathways for four ongoing research. For example, one direction would be to study bacteria in a more hydrated environment, similar to their natural habitat. Although, electrical measurements cannot be made in liquid, an environment humid enough to maintain the cells in a moist climate could restore the function of bacterial ion channels, while being conducive to CAFM, thereby allowing for the measurement of their charge transfer properties. Although multi-drug resistant strains of bacteria are highly relevant to the end application of this project, measuring a known conductive bacterium, such as *G. sulfurreducens*, may also help in the development and optimization CAFM techniques to be later be applied to less conductive strains. Furthermore, strains of *G. sulfurreducens* form conductive biofilms [22–23], offering the potential for charge transference and propagation to be studied in these much larger structures.

As denoted earlier, MRSA and *P. aeruginosa* strains were initially chosen for electrical measurement, because they are two common opportunist pathogens that readily cause wound infection. Wound infections represent a prime area where electrical current could be therapeutically applied as it may help to remove bacteria, while accelerating wound healing. In addition to comparing potential conductive differences between Gram-positive and Gram-negative bacteria, the motile nature of *P. aeruginosa* allows for the imaging and investigation of the electrical properties of pili and flagella. We plan to investigate this area further in future studies and to apply these techniques to the study of other interesting or relevant bacteria. For example, CAFM has direct measurement capabilities that could potentially help to elucidate how *G. sulfurreducens* pili exhibit metallic-like conductivity [24–25].

It is not entirely clear which component(s) of the bacteria studied here are responsible for conducting the charge. If peptidoglycan is acting as the charge carrier, it is somewhat surprising then that *P. aeruginosa* does not display a higher resistivity, as it is covered by an outer layer of lipopolysaccharides. This suggests that an alternate conductive structure may be responsible for conductance observed in this study. This is something that we could investigate in future studies, as we have several parameters worked out for examining external bacterial structures using ResiScope. For the measurement of peptidoglycan, we switched the substrate from gold-coated stainless steel to gold sputter-coated mica, because the high surface roughness of the original substrates could potentially have made it difficult to distinguish the peptidoglycan from the surface itself. Additionally, we determined that the gold-sputtered mica represents a much cleaner sample that is less easily contaminated prior to imaging. Although residual material is washed off the gold-coated stainless steel, the high surface roughness will retain a certain degree of contaminants from the environment that can obstruct electrical measurements. Conversely, the cleaved mica provides a pristine surface, ensuring that the only thing being measured is the purified peptidoglycan sample. Direct measurements of dried bacterial cells are seldom collected; most electrical measurements of bacteria focus on the planktonic state or individual live cells in solution [26]. Our work provides a framework for the future measurement of bacterial conductance in individual bacterial cells.

## 5. Conclusions

Our study successfully establishes a novel methodology for the imaging and analysis of the conductance of single bacterial cells from medically relevant bacteria, such as *P. aeruginosa* and MRSA. Our method is adaptable and can be modified as necessary to study the conductance of other bacterial species or biological substances. The average resistance values obtained in our study are comparable to those obtained in a prior study of *E. coli*, helping to validate our technique. Our work opens the door to further investigation of the electrical properties of pathogenic bacteria, information that will be essential in developing electroceuticals. Such approaches would be a great benefit for wound healing, as early studies suggest that the application of an electrical current can help to eliminate potential pathogens, while accelerating wound healing. However, to optimize such treatments, a better understanding of the localized electrical properties of adherent bacteria is necessary. Our work helps to establish a “toolbox” for investigating the electrical properties of an array of bacterial species.

## Acknowledgements

Financial support for this research was provided by the Natural Sciences and Engineering Council of Canada (Grant#400705) and the Ontario Ministry of Research and Innovation (Grant #300142). The authors would like to thank Dr. Scott Weese and Ms Joyce Rousseau of the Pathobiology Department of the University of Guelph for providing bacterial strains as clinical samples from canine patients.

